# Identification of active transposable element candidates from ROH in a de novo assembled chromosome-scale genome of a Nishikigoi, an ornamental fish derived from Common carp (*Cyprinus carpio*)

**DOI:** 10.1101/2023.12.26.573356

**Authors:** Aoi Hosaka

## Abstract

Transposable Elements (TEs) are major components of the genome. To understand their function and evolution, it is necessary to identify active TEs from a diverse range of organisms. Here, I report the genome of the Nishikigoi, an ornamental fish derived from the Common carp, and the novel approach to detecting active TE candidates.

I constructed a chromosome-scale assembly using long-read sequencing and Hi-C methods. It revealed that Nishikigoi has Robertsonian-like chromosomal translocations not seen in Common carp. I also found that Nishikigoi has a significantly different genetic background from Common carp, reflecting the intensive breeding history.

Furthermore, by focusing on Runs of Homozygosity (ROH) islands in the Nishikigoi genome and analyzing structural variations with long-read sequencing, I identified several active TE candidates.

This study not only revealed the unique genetic features of Nishikigoi but also demonstrated the potential for a novel approach in the search for active TEs.

## Main

Transposable elements (TEs), also known as “jumping genes”^1^, are major components of the genome. TEs are classified into two major classes. Class I type TEs transpose by reverse transcription of their own RNA prior to integration (copy-and-paste), whereas Class II type TEs excise their own DNA sequences to transpose (cut-and-paste). TEs, while potentially causing deleterious effects by disrupting genes through their transposition, are also recognized for significantly contributing to the genome function, diversity, and evolution^2^. Unraveling the biochemical characteristics of TEs is crucial for understanding genome evolution. However, research on the biochemical activity of transposons has been limited to those derived from a small number of species^3^. This limitation arises from the difficulty in systematically detecting active TEs from genomic information. A primary obstacle is the heterogeneity and complexity of genomes. Generally, active TEs are identified through genomic sequencing by observing structural variations^4^. However, these factors affect the accuracy in distinguishing whether these variations are caused by active TEs, allelic variations, or artifacts from the repetitive nature of TEs.

## Introduction

Runs of homozygosity (ROH) represent the genomic regions of uninterrupted long runs of homozygous genotypes, which were originally identified in humans^5^. Population history can be inferred by evaluating the frequency, size, and distribution of ROH. Fewer and shorter ROH indicate large populations, whereas more and longer ROH indicate isolated or bottlenecked populations. Since evaluating ROH is also useful for estimating the inbreeding rate, it is applied to livestock breeding researches^6^.

Nishikigoi was originally isolated from a Magoi strain of Common carp (*Cyprinus carpio*) in Niigata Prefecture, Japan around the late 17th century. Since then, intensive breeding created many strains with various colors and patterns. Nishikigoi is now referred to as “swimming jewel” and its artistic value is getting popular all over the world. According to the Agriculture, Forestry, and Fisheries Ministry trade statistics, the export value of Nishikigoi was the highest in 2022 (63 million yen), having doubled from 2013. In 2022, The Japanese government designated Nishikigoi as an important export item.

The skin color and pattern diversity of Nishikigoi are highly attractive not only for their commercial value but also for the study underlying complex genetic and environmental effects^7^. The black color of Common carp is formed by the accumulation of melanin pigment. The alternation of the pigmentation regulatory genes is suggested as one of the reasons for the color pattern diversities. *Melanocrtin cereptor 1* (mc1r) is a G protein Coupled Receptor (GPCR) mediating the physiological action of melanocortins by activating the cyclic AMP (cAMP) activating pathway^8^. Indeed, expression levels of the mc1r gene vary among Nishikigoi strains^9^. The expression level and sensitivity of the Melanin-concentrating hormone 1 (mch1), which pales the skin color by stimulating the aggregation of pigment, is reduced in Tasho-Sanshoku, a variety of black, white, and red-colored strain, compared to Common carp. This phenotype might have been selected due to the robustness of the dark skin color against environmental changes. In addition to the protein-coding genes, microRNAs and long non-coding RNAs are involved in pigmentation^10‒13^.

Nevertheless, neither the responsible genes for the breeding targets nor the impact of the breeding processes on genetic variations is largely unknown due to the lack of Nishikigoi genomic information. Nishikigoi’s genetic background is highly complex since various strains from all over the world, such as “Doitu-Goi,” a German strain, were crossed to create novel patterns. In addition, Nishikigoi have been experienced extremely strong selection pressures through generations; from tens of thousands of offspring, only a few individuals with the most valuable traits are selected for breeding each generation. Notably, Ojima et al. found that the karyotype of Nishikigoi differs from Common carp ^14^. While Common carp has 100 paired chromosomes (2n=100), Nishikigoi strains have several unpaired chromosomes, long chromosomes, and micro-chromosomes. Taken together, Nishikigoi genome would be highly differentiated from Common carp. Therefore, to study the genetics of Nishikigoi, it is necessary to first decipher the genome.

In this study, I report the chromosome-scale assembly of Nishikigoi strain, GinrinShowa. The Hi-C scaffolding method revealed that the aneuploidy in Nishikigoi was caused by chromosomal translocations. In addition, I detected active TE candidates using long-read sequencing with focusing on ROH islands in a Nishikigoi strain. Utilizing a newly determined genome to search for active TEs offers a broadly applicable method and is expected to contribute to a more general understanding of TEs.

## Materials and methods

### Materials

A GinrinShowa and a Koganeryu of Nishikigoi were bought from Maruhiro Koi Farm (Niigata, Japan) and Maruju Koi Farm (Niigata, Japan), respectively. They were grown in Prof. Norichika Ogata’s private pond (Kanagawa, Japan) (**Supplementary Figure S1A, C, and D**).

### DNA extraction and genome sequencing

The DNA was extracted from gills of a GinrinShowa and a Koganeryu using NucleoBond HMW DNA (Takara) following the manufactures’ protocol.

For short-read sequencing, MGIEasy FS DNA Library Prep Set (MGI) used for the library preparation. The prepared libraries were sequencing by DNBSEQ (MGI) at 150bp paired-end mode. The library preparation and sequencing were performed at GenomeLead Inc. For long-read sequencing, High-molecular-weight DNA was enriched by BluePippin (sage science), and the DNA library was prepared with the SMRTbell Express Template Prep Kit 2.0 (PacBio). The prepared library was sequenced by Revio (PacBio). The library preparation and sequencing were performed at Macrogen Japan Inc. The obtained HiFi reads were used for the downstream analyses.

### Hi-C library preparation

The Hi-C library was prepared as described previously ^15^ with several modifications. About 300mg of frozen muscle was ground with Liquid nitrogen. To crosslink, the sample was thawed in 1ml of fixation buffer (1x PBS buffer with 1% formaldehyde) and incubated for 10 minutes with gentle mixing. After the filtration with 200 µm cell strainer, the sample was pelleted 3 times (3,000 x g for 5min) and resuspended with 1 x PBS buffer. The Washed sample was resuspended by 1ml of 1x PBS buffer. DNA concentration was measured to determine the input DNA for Hi-C reactions.

The nuclei containing about 1,000 ng of DNA were resuspended with 1x M buffer (Takara). To permeabilize nuclei, the nuclei were pelleted by centrifugation (3,000 x g for 5min), resuspended with 100 µl of 0.5 % SDS, and incubated at 65 ℃ for 5 minutes. 120 µl of 2% Triton was added and incubated 37 ℃ for 15 minutes. The DNA was digested by 10 µl of Hind III (Takara) at 37℃ overnight in 250 µl of 1x M buffer solution. The nuclei were pelleted centrifugation (1,500 x g for 5min) and resuspended with Fill-in mastermix (0.4 mM biotin-14-dATP, 1.5 µl of 10 mM dCTP, 1.5 µl of 10 mM dGTP, 1.5 µl of 10 mM dTTP, 20 µl of NEBuffer 2, 134 µl of water). 3.5 µl of Klenow Fragment (NEB) was added and incubated at 37℃ for 2 hours.

The nuclei were pelleted by centrifugation (1,500 x g for 5min), resuspended with T4 DNA ligation mastermix (40 µl of T4 DNA Ligase Reaction buffer (NEB), 10% Triton X-100, 20 µl of T4 DNA Ligase) and incubated at 25℃ for 4 hours.

The nuclei were pelleted by centrifugation (1,500 x g for 5min), resuspended with 500 µl of SDS Lysis buffer (50 mM Tris-HCl, 10 mM EDTA, and 1% SDS), 8 µl of 8 M NaCl. 4 µl of RNase A (100 mg/ml; Thermo) was added and incubated at 37℃ for 30 minutes. To reverse-crosslink, 4 ul of Proteinase K (20 mg/ml; Invitrogen) was added and incubated at 65℃ overnight. 500ul of Phenol/Chloroform/Isoamylalcohol was added and mixed vigorously. The sample was centrifuged at 10,000 x g for 5 minutes and the supernatant was transferred to the new microtube. 50 µl of 3M NaOAc and 450 µl of Iso-propanol were added and mixed. The DNA was precipitated by centrifugation (13,000 x g for 20min at 4℃). The DNA was rinsed by 500 µl of 70% Et-OH and eluted by 50 µl of Tris buffer. The DNA concentration was quantified by Qubit 3.0 (Thermo).

500 ng of ligated DNA was sheared into approximately 500 bp by Covaris M220 instrument in a 130 µl microtube. 5 µl of Dynabeads MyOne Streptavidine C1 beads were washed once by TWB buffer (5mM Tris-HCl pH 8.0, 0.5mM EDTA, 1M NaCl, 0.05% Tween-20). The beads were captured by a magnetic stand and the supernatant was removed. The beads were resuspended by µl of 130 2x BB buffer (10mM Tris-HCl pH 8.0, 1mM EDTA, 2M NaCl), mixed with the sheared DNA, and incubated for 15 minutes with gentle mixing. The beads were washed with 500 µl of TWB buffer twice, 200 µl of water once. Then the beads were resuspended with 12.5 µl of water.

1.75 µl of End Repair & A-tailing Buffer (KAPA) and 0.75 µl of End Repair & A-tailing Enzyme Mix were added to the beads and incubated at 20℃ for 30 minutes, and then 65℃ for 30 minutes. The reaction solution was mixed with 7.5 µl of Ligation Buffer, 2.5 µl of DNA Ligase, and 1.25 µl of Dual-index adapter (diluted 10 times) and incubated at 20 ℃ for 15 min. The beads were washed twice with 500 µl of LWB, once with 500 µl of Tris buffer, and resuspended with 5 µl of Tris buffer. Then, 6.25 µl of 2× KAPA HiFi Hot Start Ready Mix and 1.25 µl of 2x Library Amplification Primer Mix were added to the solution, and polymerase chain reaction (PCR) was performed with following conditions; 98 ℃ for 45 sec, then, 10 cycles of 98 ℃ for 15 sec, 60 ℃ for 30 sec, and 72 ℃ for 30 sec, and a final extension at 72 ℃ for 1 min. The supernatant of the PCR product was purified and dual size-selected using 0.5x and 0.3x volumes of AMPure XP beads (Beckman) and eluted with 13 µl of TE buffer. The library was qualified, then sequenced by Novaseq system (illumina) at Macrogen Japan Inc.

### Genome assembly and scaffolding

The HiFi reads and Hi-C reads were used for the *de novo* assembly by Hifiasm 0.19.6-r595^16^. “--hg-size 1.7g --hom-cov 54” options were specified based on previously reported genome size and the obtained data amount. The obtained primary contigs were used for scaffolding. The reads were aligned to the contigs using Juicer 1.6^17^. The generated contact maps were then used for scaffolding with a 3D-DNA pipeline^18^ with following options “--rounds 0 -- editor-coarse-resolution 2500000 --editor-coarse-region 12500000 --editor-coarse-stringency 1 --polisher-coarse-resolution 2500000 --polisher-coarse-region 15000000 -- polisher-coarse-stringency 1 --splitter-coarse-resolution 2500000 --splitter-coarse-region 15000000 --splitter-coarse-stringency 1” to reduce unwanted error corrections. The scaffolds were reviewed and manually corrected using JuiceBox 1.11.08 (https://github.com/aidenlab/Juicebox). The final assembly was named Sh_v1.0 and used the following analyses.

### Assessment of genome assembly

Contiguity of the assembled contigs and the assembly are evaluated by Seqkit ^19^ (2.5.1). Completeness of the assembly was assessed by Benchmarking Universal Single-Copy Orthologs (BUSCO)^20^ (5.5.0) against Actinopterygii database.

### Annotation

Protein-coding regions were predicted by aligning the genes from Common carp^21^ using LiftOff^22^ (v1.6.3). Transposable elements were predicted by EDTA^23^ (v2.0.1). To filter out gene-related sequences, predicted CDS sequences were provided. Densities of genes and each transposon family in each 1 Mb were calculated by BEDtools^24^. Distribution of the genes and the transposable elements were visualized by Circos^25^. Correlation efficiency of the distribution of the genes and the each transposon family in each 1 Mb bin was calculated and visualized as heatmap using R Complexheatmap^26^ package. To estimate Kimura substitution level of the TEs, TEs were re-annotated with RepeatMasker (4.1.2-p1) using the TE library generated by EDTA^23^ as reference sequences. Kimura substitution level was calculated using a RepeatMasker utility script “calcDivergenceFromAlign.pl”. The result was visualized by R after excluding minor TE families.

### Detection of Collinearity regions

Detected coding regions were then used for the detection of collinearity regions by the Python version MCScan^27^ (1.3.8). The homologous gene blocks, which contain more than 30 gene pairs, were defined as collinearity regions, and used for drawing synteny plots.

### Resequencing

The whole-genome sequencing datasets of two Nishikigoi strains GinrinShowa, Koganeryu, wild Common carp “Songpu” PRJNA684795 and “Yellow River” PRJNA684676, and a domesticated strain “Furui” PRJNA684797 were used to estimate the genetic variations. The reads were aligned to the assembly by bowtie2^28^ (2.5.1) with default parameters, and the resulting SAM files were converted to sorted BAM files by SAMtools^29^ (1.18). The coverages in each 10kb bin were calculated with mosdepth^30^ (0.3.3). bcftools (1.17) was used for call variants with diploid mode. The resulting vcf files were further filtered using VcfFilter (https://biopet.github.io/vcffilter/) to extract DP > 5 and MQ > 30 sites. Principal Component Analysis was computed using the filtered sites by PLINK^31^ (1.9). To define Runs Of Homozygosity (ROH) regions, firstly, the number of heterozygous sites in each 10kb bin was counted by bedtools^24^ (2.31.0). The regions fewer than 10 heterozygous sites in a 10 kb bin lasting more than 500kb were defined as ROHs. The ROHs were visualized on the IGV^32^ (2.11.9) genome browser.

Fluctuations of effective population size through generations were estimated by Pairwise Sequentially Markovian Coalescent (PSMC) model^33^. First, I obtained the whole-genome consensus sequences by SAMtools (1.9) and bcftools (1.9) with following command “mpileup -C50”, then “bcftools call -c - | vcfutils.pl vcf2fq -d 10 -D 100 | pigz”. The resulting fq.gz file were converted into psmcfa format by fq2psmcfa command with a “-q 20” option. To infer population size history, psmc command was used with “-N25 -t15 -r5 -p "4+25*2+4+6" options. The result was summarized with psmc_plot.pl command with “-g 5 -R” options. The generation time was assumed as 5 years. The samples that encountered issues and failed to produce results, as well as samples indicating abnormal values, were excluded from the analysis.

### Detection of active TE candidates in ROH islands

HiFi reads were aligned to the assembly using pbmm2 (1.13.0) with HiFi mode (https://github.com/PacificBiosciences/pbmm2), and structural variations were called using pbsv (2.9.0) (https://github.com/PacificBiosciences/pbmm2) with default parameters. From the resulting vcf file, the hemizygous insert or deletion sites (InDels) with more than 500bp, which located in ROHs were extracted. The sequences of extracted InDels were used for pairwise homology search with blastn^34^. The sequences satisfying the following criteria were extracted and grouped: (1) The sequence length difference between query and subject is smaller than 100 bp, (2) The homologous region covers more than 90 % of subject sequences, (3) More than 3 sites are found that satisfy the (1) and (2) criteria. The multiple alignment of the sequences of the grouped InDels was generated by mafft with a “--adjustdirectionaccurately” option. Then, the consensus sequences of the grouped InDels were constructed using EMBOSS cons command (6.6.0.0) from the result of the multiple alignment (https://emboss.sourceforge.net/apps/cvs/emboss/apps/cons.html). TE sequences in the consensus sequences were defined using RepeatMasker (4.1.2-p1) with the TE library generated by EDTA^23^.

## Results and Discussion

### Genome Assembly

To assemble a Nishikigoi genome, DNA from a GinrinShowa variety was sequenced by Revio sequencer. 92.4 Gb of HiFi reads with N50 14.4 kb were obtained (**Supplementary Table 1**). All reads were used for *de novo* assembly with hifiasm. I obtained 385 primary contig with total length 1,58 Gb and with N50 25.3 Mb. To scaffold the contigs, I constructed a Hi-C library with Hind III restriction enzyme from a same individual and sequenced on NovaSeq X sequencer. 261 M read pairs were obtained and used for scaffolding. As a result, I obtained 200 scaffolds with total length 1,57 Gb and with N50 30.5 Mb (**Figure. 1A**). The previously reported Common carp genome (GCF_018340385.1) consists of 6,701 scaffolds from 19,838 contigs, indicating that my assembly is much more contiguous. I assessed the BUSCO score of the assembly against Actinopterygii database to further validate the completeness and the accuracy of the assembly. 3,694 genes (98.8 %) out of 3,640 genes were detected. The Complete BUSCOs were slightly higher than the previous assembly (97.8 %) confirming that my assembly is superior to the previous assembly (**Supplementary Table S2**). Substantial number of genes are defined as Complete and duplicated BUSCOs, reflecting the allotetraploid nature. I named the assembly as Sh_v1.0 and used it as the reference genome following analyses.

**Fig. 1.**
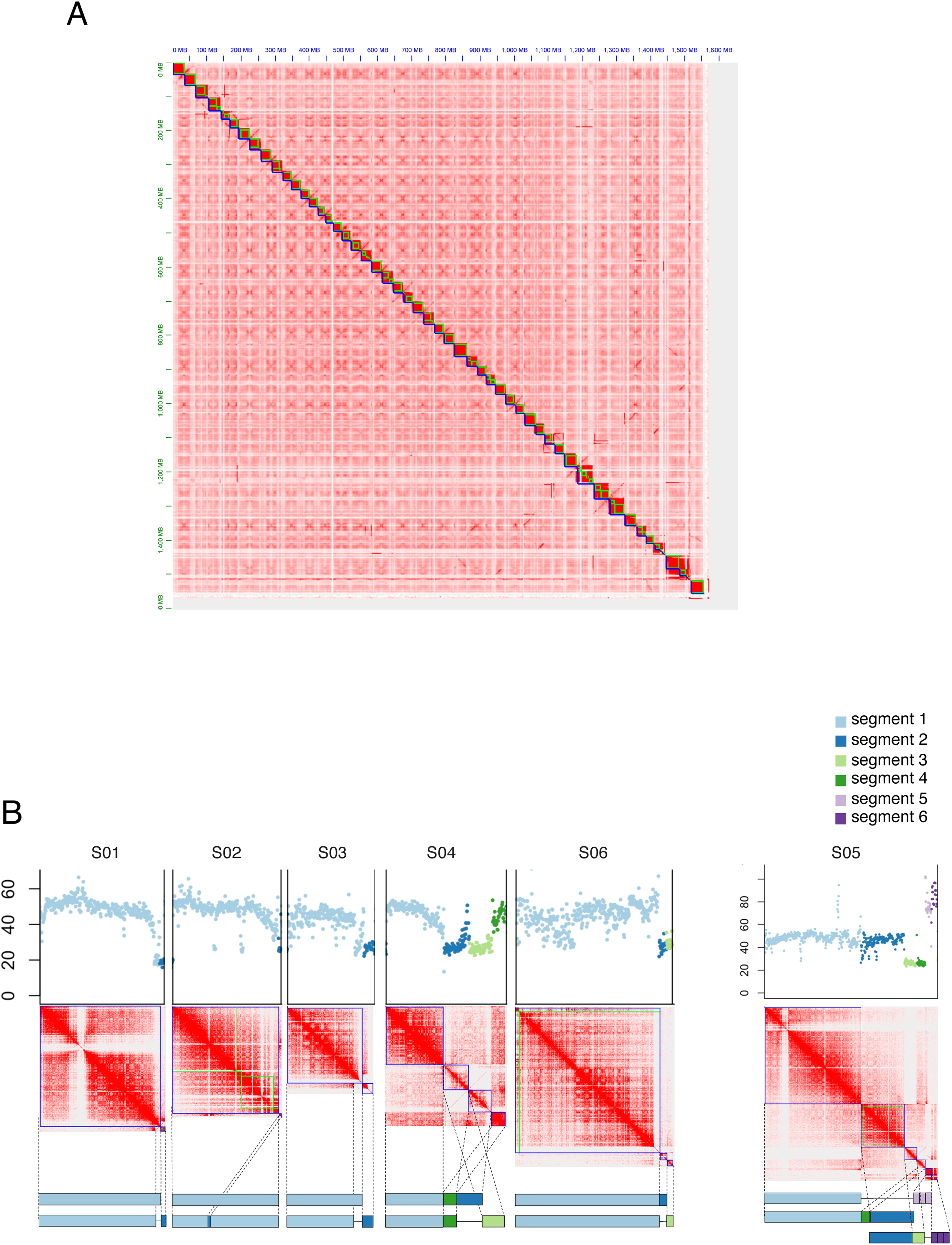
Chromosomal assembly of a Nishikigoi. (A) Hi-C heatmap of the assembly. Green and blue lines indicate contigs and scaffolds, respectively. (B) Putative chromosomal structures of unresolved contigs. Upper panels indicate the coverages in each 10 bins. Each contig was colored differently. Middle panels are Hi-C heatmaps of grouped contigs based on the contact intensity. Lower panels indicate putative chromosomal structures based on the coverages in each contig and Hi-C contact intensity among contigs.

Among the scaffolds, 43 scaffolds formed chromosomes and 137 scaffolds, which were much shorter than chromosomes, were unanchored (Supplementary Table S3, S4). The remaining 20 scaffolds could not be anchored because these scaffolds showed complex contact patterns with other scaffolds (**Figure 1B** **middle panel**). These scaffolds were divided into 6 groups based on the contact patterns. I referred to these scaffolds as chromosomal segments. In the case of S01 groups, which consist of two chromosomal segments, the subterminal region of chromosomal segment 1 interacts with chromosomal segments, while terminal regions of chromosomal segment 1 do not based on the Hi-C heatmap.

To understand the precise structure of the chromosomal segments, I checked the sequencing coverage on the chromosomal segments by aligning 273 M read pairs of whole-genome short-read sequencing data of a GinrinShowa. I noticed that whereas the mean coverage in all regions was 47.29, the coverages at the terminal region of chromosomal segment 1, and chromosomal segment 2 were nearly half to mean coverage (**Figure 1B** **upper panel**). This indicates that these regions exist only in either chromosome in a diploid chromosome pair. The possible chromosome structure was visualized in the lower panel of **Figure. 1B**. Similar patterns were observed in S02 and S03 groups, except chromosomal segment 2 of group S02 interacts with the internal region of chromosomal segment 1. The group S04 and S06 were also able to predict possible chromosome structures based on the same approach (**Figure 1B** **lower panel**).

However, in the case of the S05 group, while chromosomal segment 1 interacts with chromosomal segment 2, 4, and 5, Chromosomal segment 2 interact with 3,4, and 6. These interaction patterns cannot be explained unless hypothesizing aneuploid chromosomes, as visualized in the lower panel of **Figure 1B**. Notably, this hypothesis is consistent with karyotyping results as Ojima et al. reported^14^. Therefore, it is highly probable that the aneuploidy is the important chromosomal feature of Nishikigoi.

### Annotation

To compare the syntenic structure of the assembly to Common carp genome, I predicted genic regions by homology-based annotation with LiftOff^22^ using Common carp gene structures as queries. As a result, 48,925 genes were predicted. 43 chromosomes and 5 out of 6 groups of chromosomal segments of the assembly were highly syntenic with the Common carp genome (**Figure 2A**). Importantly, In the group S05, whereas chromosomal segment 1 was syntenic to A05 chromosome, chromosomal segment 2 was syntenic to A19 chromosome (**Figure 2B**). The aneuploidy in Nishikigoi was possibly caused by the translocation between A05 and A19 chromosomes, which is similar to Robertsonian translocation^35^. Aneuploidy is one of the main reasons of sterility, embryonic lethality and severe developmental disorders^36^. Nishikigoi strains are highly fertile and crossable to various strains including wild Common carp populations. It would be interesting to study how the aneuploid chromosomes behave during reproduction for normal fertilization.

**Fig. 2.**
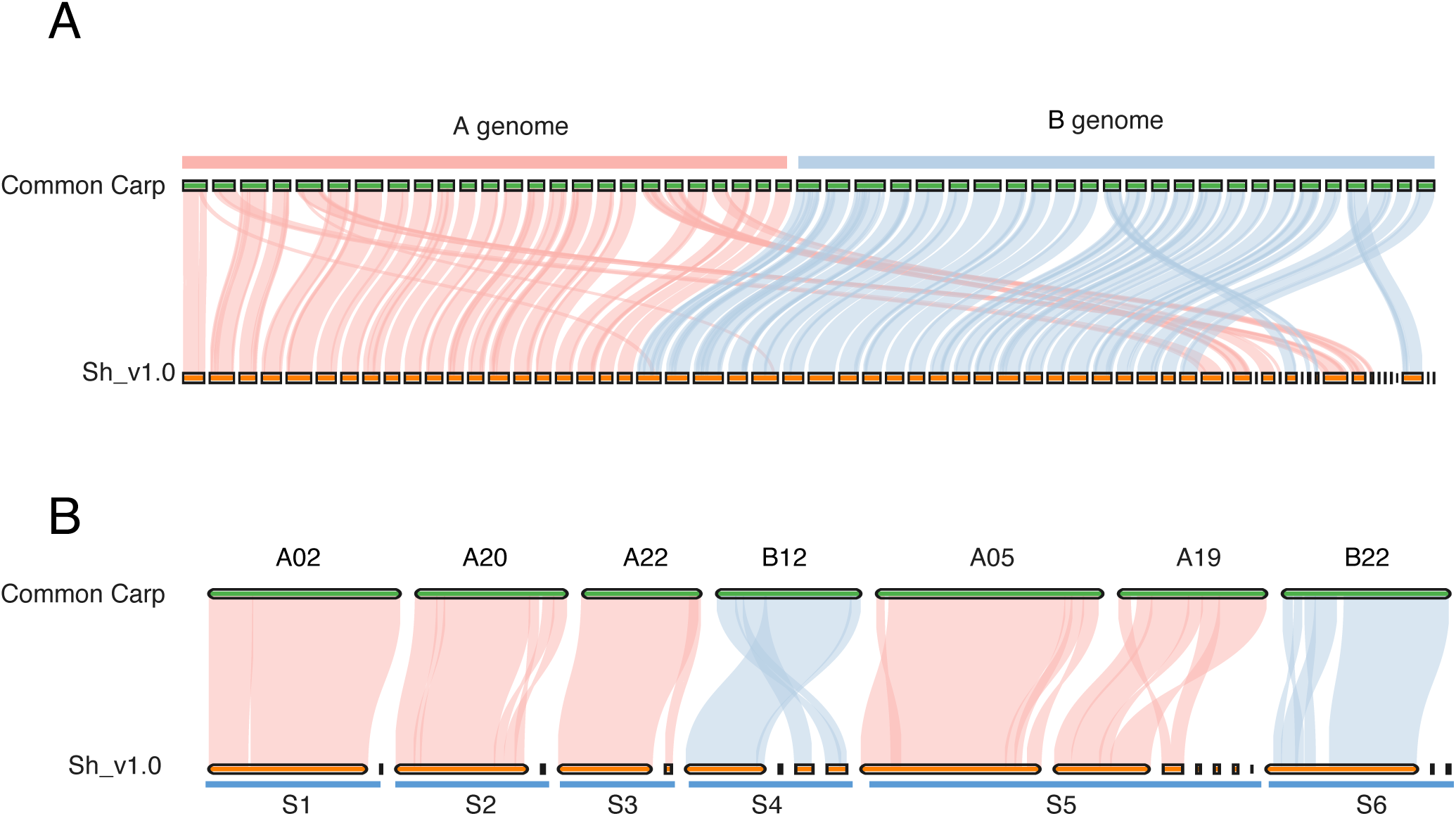
Syntenies between common carp and Nishikigoi. (A) Synteny plots between common carp and Nishikigoi. Red and blue lines indicate syntenic regions between A genome and B genome of common carp, respectively. (B) Synteny plots focused on Chromosomal segments in Nishikigoi.

### Breeding signatures in Nishikigoi individuals

To investigate the phylogenomic relationship between Nishikigoi and Common carp populations, I use the whole-genome sequencing data of two Nishikigoi strains, named Ginrinshowa and Koganeryu, and publicly available 3 Common carp populations, named Songpu, Furui, and Yellow River. Songpu and Yellow River strains are a wild population sampled at Beijing, Wuxi (Jiang Su Province), respectively. Whereas Furui strain is domesticated populations sampled at Dong Er (Shandong Province). These sequence datasets were mapped to the Sh_v1.0 genome. On average, 92.1% of reads were mapped (Supplementary Table S5), and 61,230,274 polymorphic sites were called. A Principal Component Analysis (PCA) and distance matrix analysis supported previous results, in which the Common carp strains were classified into two major clades; the European clades (Songpu strain) and Asian Clades (Furui and Yellow River strain) (**Figure 3A, B**). Importantly, the two Nishikigoi individuals have highly different genetic backgrounds from these Common carp strains and even with Nishikigoi individuals. Next, I inferred the population size history of these populations using pairwise sequential Markov coalescent (PSMC). The effective population size of Songpu strain has been declining after reaching its maximum around one million years ago (Supplementary Figure S2). On the other hand, the effective population sizes of Furui, Yellow River, and two Nishikigoi strains also showed more recent (around 0.1 million years ago) and dramatic expansion. It should be noted that differences in library preparation methods and sequence coverages would affect the result^37^, direct comparison of Nishikigoi strains and these Common carp strains poses difficulties.

**Fig. 3.**
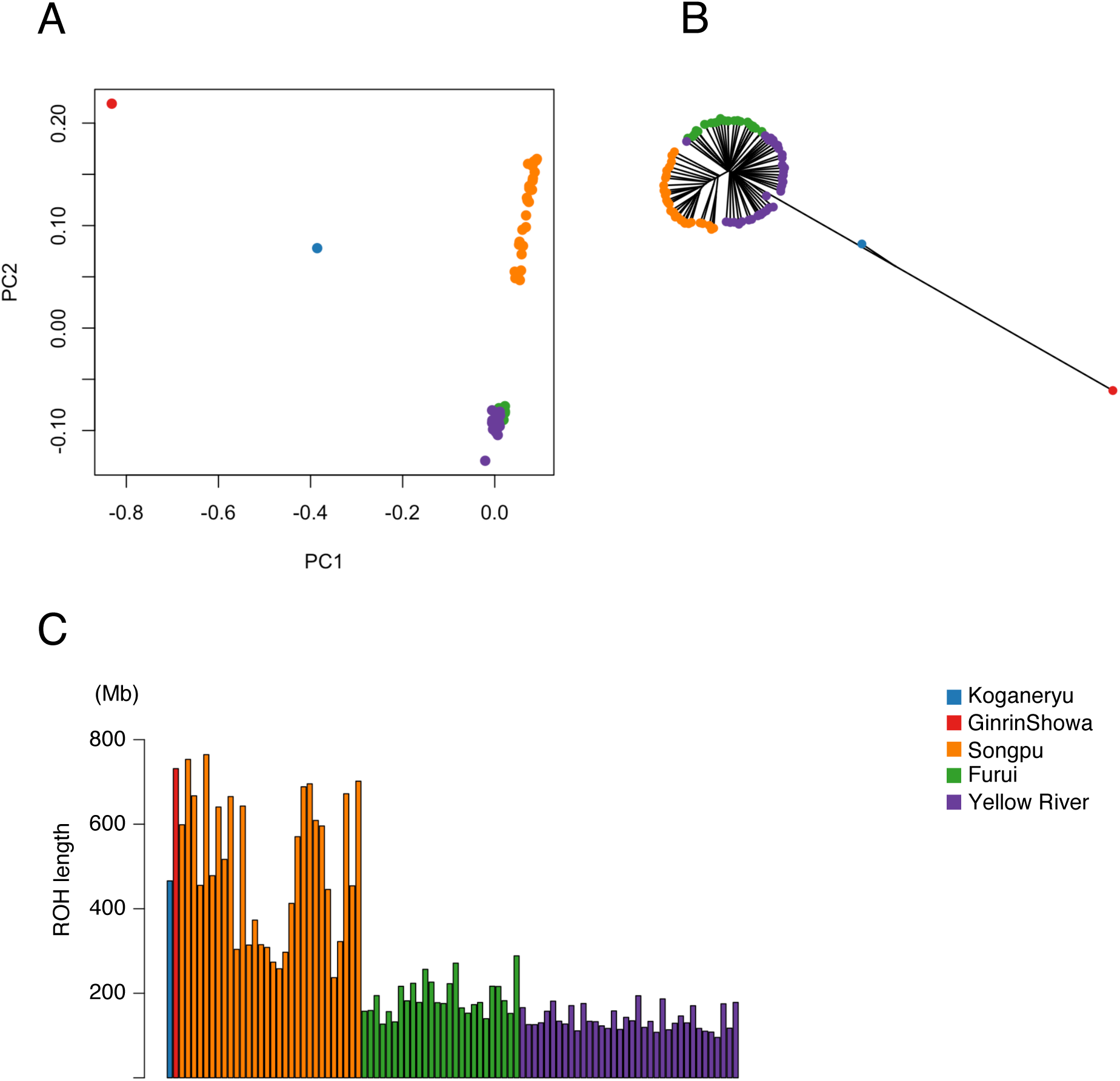
Genetic variations of Common carp strains and Nishikigoi strains. (A) PCA plot showing the genetic structure of 91 Common carp individuals from 3 strains and 2 Nishikigoi strains. (B) Neighbor-joining tree based on a distance matrix. (C) Bar graph showing the sum of the ROH regions. Different colors indicate separate strains as shown in the legend.

To further investigate the breeding history, I searched ROH islands. Interestingly, the length of ROH islands in Nishikigoi strains are 2 to 4 times longer than the mean ROH length of domesticated carp Furui, Indicating the occurrence of a strong bottleneck during Nihikigoi breeding (**Figure 3C**). I also

To create strains with novel colors and patterns, Nishikigoi breeders have extensively crossed various strains. Once elite strains are established, these strains are inbred to maintain the characters. High genetic diversity with long ROH islands in Nishigoi strains observed in the above results may reflect such Nishikigoi’s breeding history.

### Detection of active TE candidates

TEs were annotated using EDTA pipeline^23^. In total, 38.42 % of the genome was covered with TEs, and hAT type TEs were most abundant (11.34 %) (Supplementary Table S6). Each TE family shows distinct distribution pattern. Whereas Helitron, CACTA and Tc1 Mariner TE families tends to locate close to gene-rich regions, hAT, Copia, Gypsy and Mutator Families tends to locate gene-poor regions (**Figure 4A, 4B, Supplementary Figure S3**). Next, I estimated the sequence divergence rate of the identified TEs by calculating Kimura 2-parameter distances. This value reflects with TE ages. Notably, TEs sharing highly similar sequences were most abundant, indicating these TEs have proliferated recently or may still have transposition activity (**Figure 4C**).

**Figure 4.**
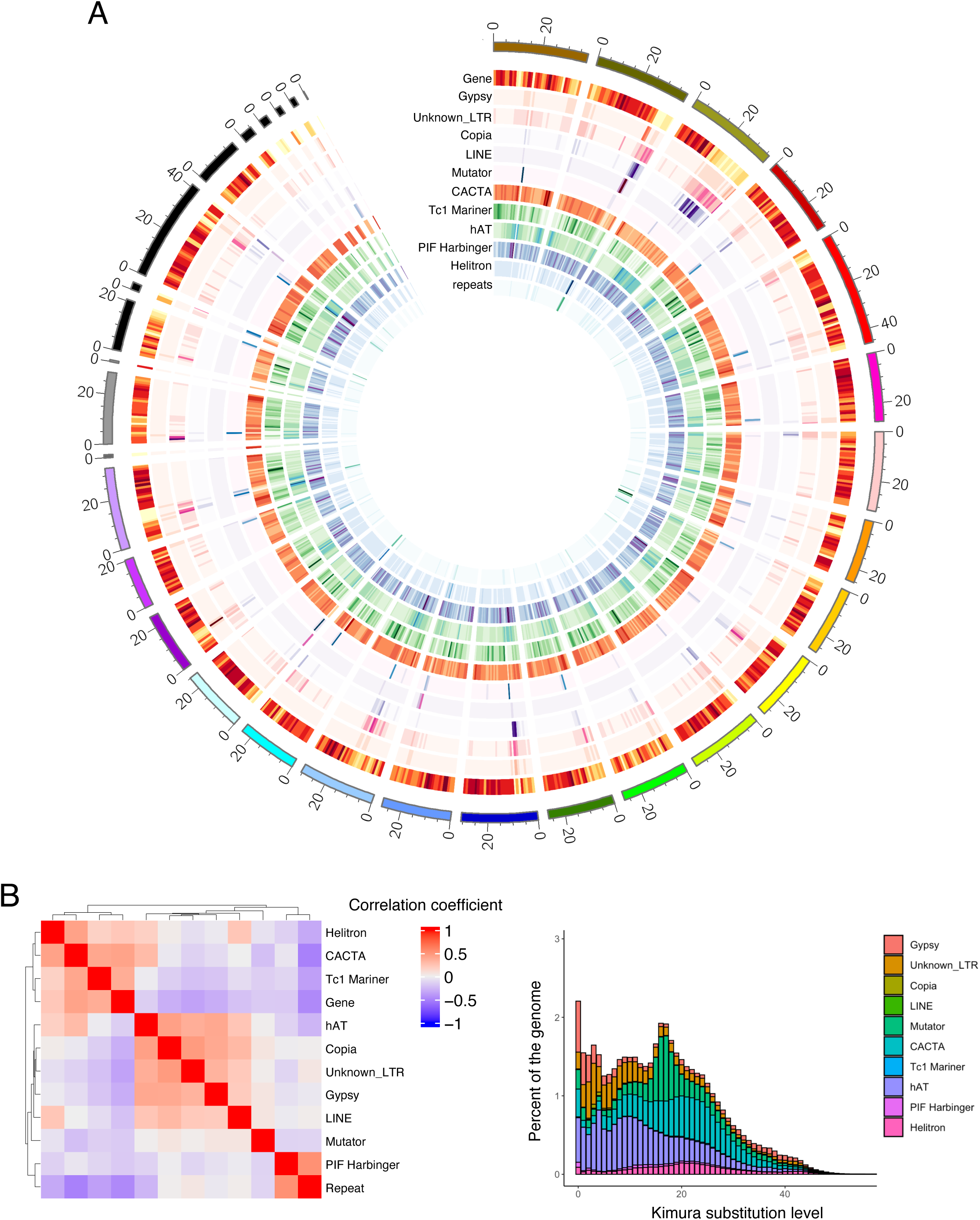
Distribution of TEs in the Nishikigoi genome. (A) Chromosomal distribution of genes, and transposons in A-genome of Nishikigoi genome. Heatmap indicates the density of the genes and the TE families in each 1Mb bin. (B) Heatmap showing the correlations of their locations. (C) The bar graph represents the genome coverage of each TE family classified by the Kimura substitution levels. Y-axis and X-axis indicate genome coverage of each TE (%) and Kimura substitution levels, respectively.

**Table 1.** Assembly Statistics of GinrinShowa and common carp.

|  | Primary Contig | Final Assembly (Sh_v1.0) | Common carp GCF_018340385.1 |
| --- | --- | --- | --- |
| Contigs | 385 | 200 | 6,701 |
| Total length (bp) | 1,580,504,010 | 1,576,735,516 | 1,680,134,903 |
| Mean length (bp) | 4,105,205.20 | 7,883,677.60 | 250,729 |
| N50 (bp) | 25,303,196 | 30,506,629 | 29,545,497 |

Since the very recent insertions of TEs are hemizygous, I hypothesized that by analyzing sequences displaying hemizygous insertions or deletions detected within Runs of Homozygosity (ROH) regions, it might be possible to explore active TE candidates (**Figure 5A**).

**Figure 5.**
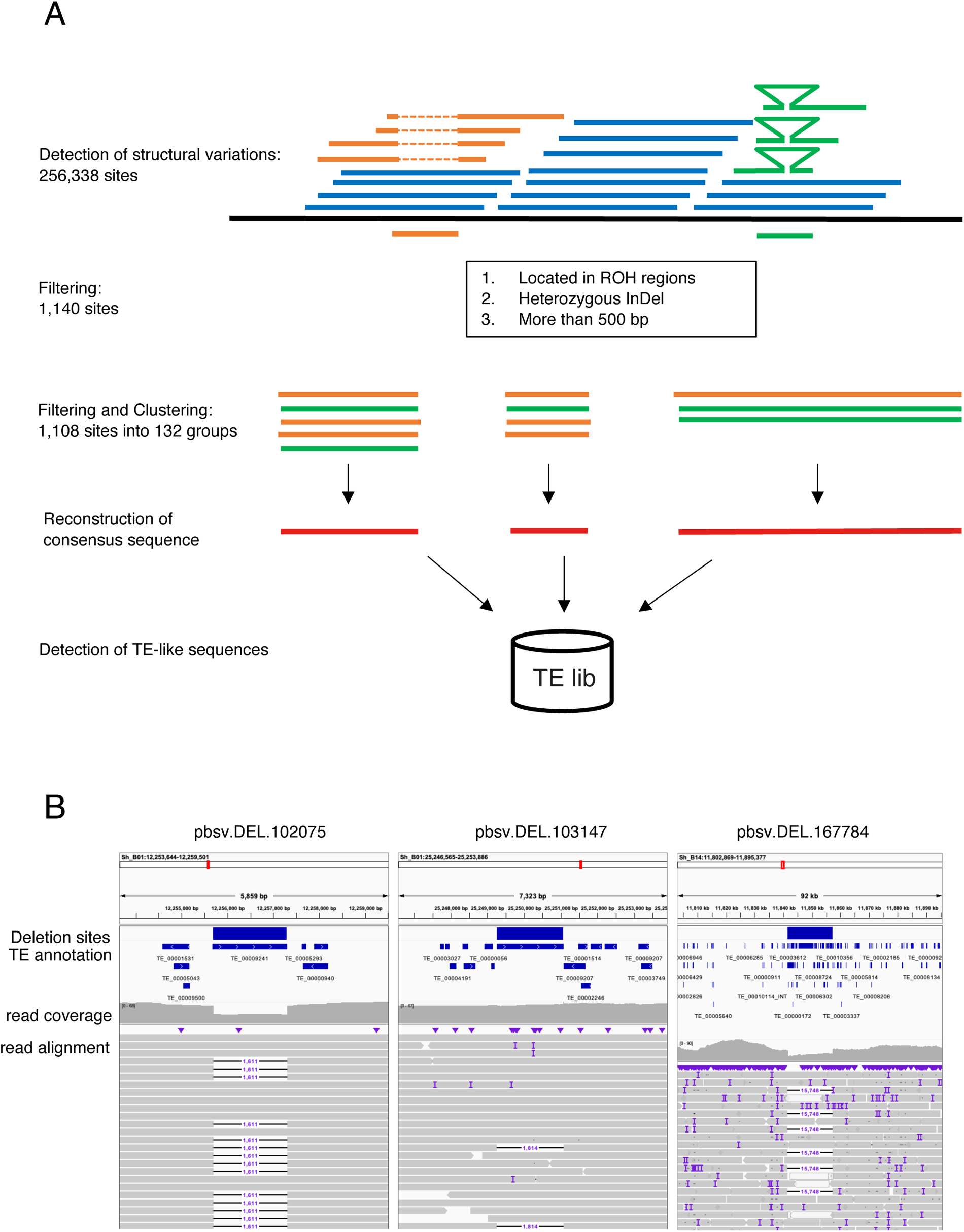
Detection of active TE candidates. (A) A schematic diagram showing the analytical pipeline to detect hemizygous INDEL and search active TE candidates. (B) Typical examples of heterozygotic deletions (left: pbsv.DEL.102075, middle: pbsv.DEL.103147, and right: pbsv.DEL.167784) possibly caused by transposition of TEs. Heterozygous deletions, TE distribution, coverages of HiFi reads, and alignment patterns of HiFi reads are visualized on a genome viewer.

To test the idea, I aligned the HiFi reads, which were used for *de novo* assembly, to the assembly and searched structural variations. Among 256,338 variations, 1,140 sites were either insertions or deletions with more than 500 bp found in ROHs of GinrinShow strains. The active TEs would insert into the genome several times, and therefore such TEs would be detected several times as hemizygous InDels. Based on the expectation, I performed a pairwise homology search using the sequences of the filtered InDels to extract the sequences that are found more than 3 times in the hemizygous InDels (see Methods). 1,108 among 1,140 sites were detected and were classified into 132 groups based on their sequence length and homology (Supplementary Table S7). The numbers with the consensus sequences in each group were generated and TE-like sequences were searched. As a result, 126 consensus sequences have similarities with TEs (Supplementary Table S8). The number of InDels classified into each group varied significantly. pbsv.DEL.102075_group, which contains a hAT sequence, had the most with 106 InDels. The next highest was pbsv.DEL.103147_group with 84 InDels. The sequence length of each group was also varied significantly. The longest group was pbsv.DEL.167784_group with 15,749 bp. This sequence was composed from 38 different TEs. Indeed, the hemizygotic nature of these regions was observed at the genome browser, supporting the results (**Figure 5B**). These results suggest that both class I and class II type TEs may have transpose activity. In addition, highly chimeric TEs are may also able to transpose like Pack-MULEs^38,39^.

## Conclusion

Here, I reported a chromosome-scale assembly of Nishikigoi using HiFi reads and Hi-C technology (**Figure 1A**). This assembly is the most contiguous and accurate genome among published Common carp genomes. Both the cytological observation and Hi-C technology indicated that Nihsikigoi has aneuploid chromosomes caused by translocation (**Figure 1B, 2B)**. It would be interesting to dissect how the aneuploid chromosomes are inherited through generations.

Population analyses shed light on the complexity of Nishikigoi genetic background, Although the sample size is not enough to discuss the genetic diversity of Nishikigoi strains (**Figure 3**). The exquisite patterns of Nishikigoi are the result of breeders’ highly meticulous and strategic breeding programs, coupled with extensive selective processes. In the absence of such rigorously controlled breeding, the value of these fish can easily diminish, even within a few generations of random breeding (Supplementary Figure S1E). The genome information reported herein is likely to serve as fundamental data for the molecular genetics of Nishikigoi. I also proposed a novel method to detect active TE candidates using HiFi reads and by focusing on ROH islands (**Figure 5**). Since this method is simple and widely applicable to eukaryotic organisms, this would contribute to the progress of TE research.

## Acknowledgment

This work was partly supported by a joint research program of Yokohama City University. The Human Genome Center at the University of Tokyo provided supercomputing resources. The experiments, including all sequencing and data analyses, were funded by a private grant from Nihon Biodata Corp.

I wish to thank Prof. Ogata Norichika and Colleagues of Nihon Biodata Corp. for carefully reading and giving critical comments on this manuscript. And lastly, this paper is dedicated to GinrinShowa and Koganeryu, beloved by Prof. Ogata, with heartfelt condolences.

## Data availability

The raw DNA sequencing reads, genome assembly have been deposited into the DDBJ databank under BioProject Number PRJDB16206.

---

**Supplementary Figure S1.**
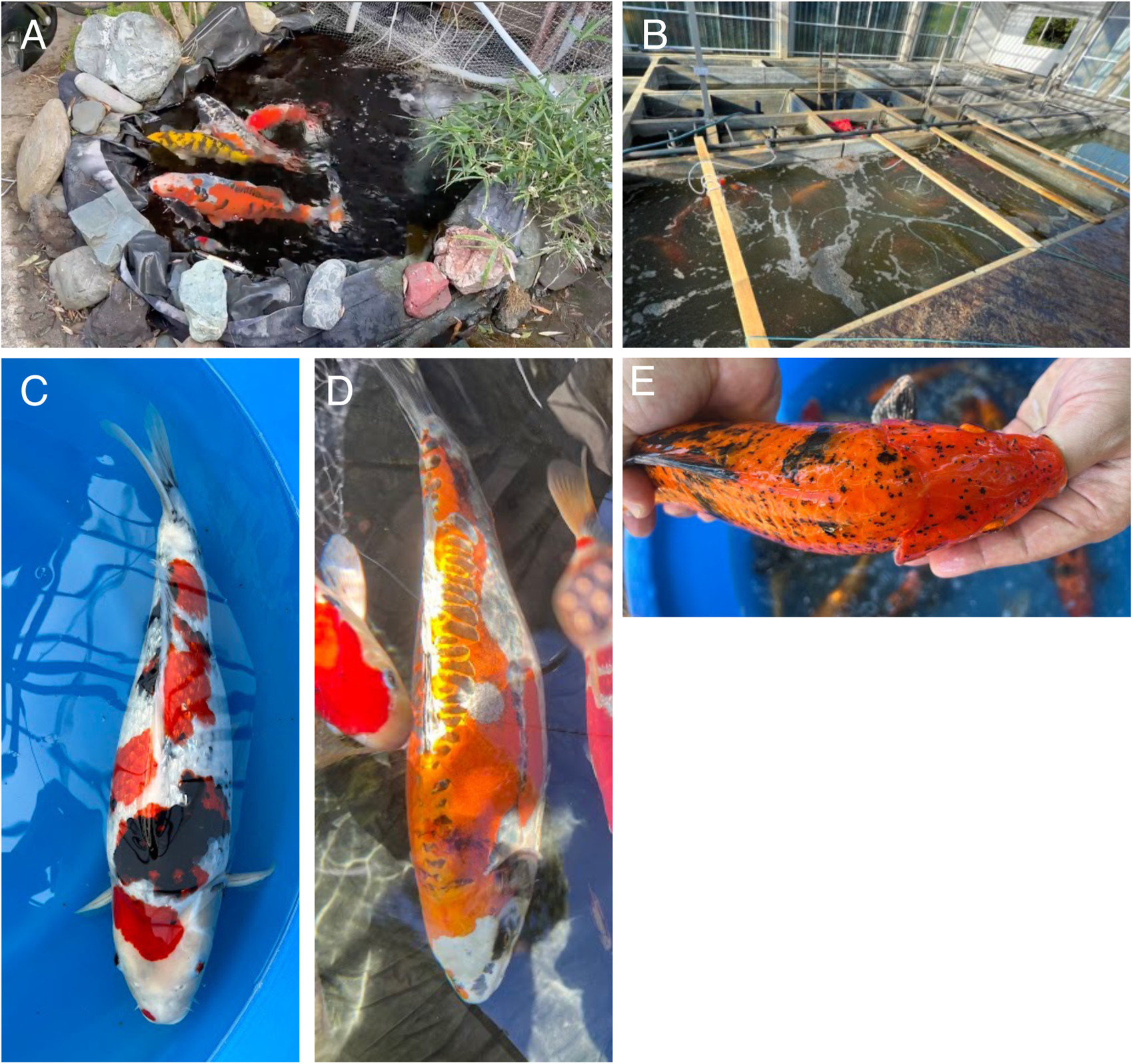
Color and pattern variations among Nishikigoi strains. Private ponds in Kasawaki, Japana (A), and Dambulla, Sri Lanka (B). GinrinShowa (C) and Koganeryu (D) used in this study grown in a private pond in Kawasaki, Japan. An individual born from random crossing, grown in Dambulla, Sri Lanka (E).

**Supplementary Figure S2.**
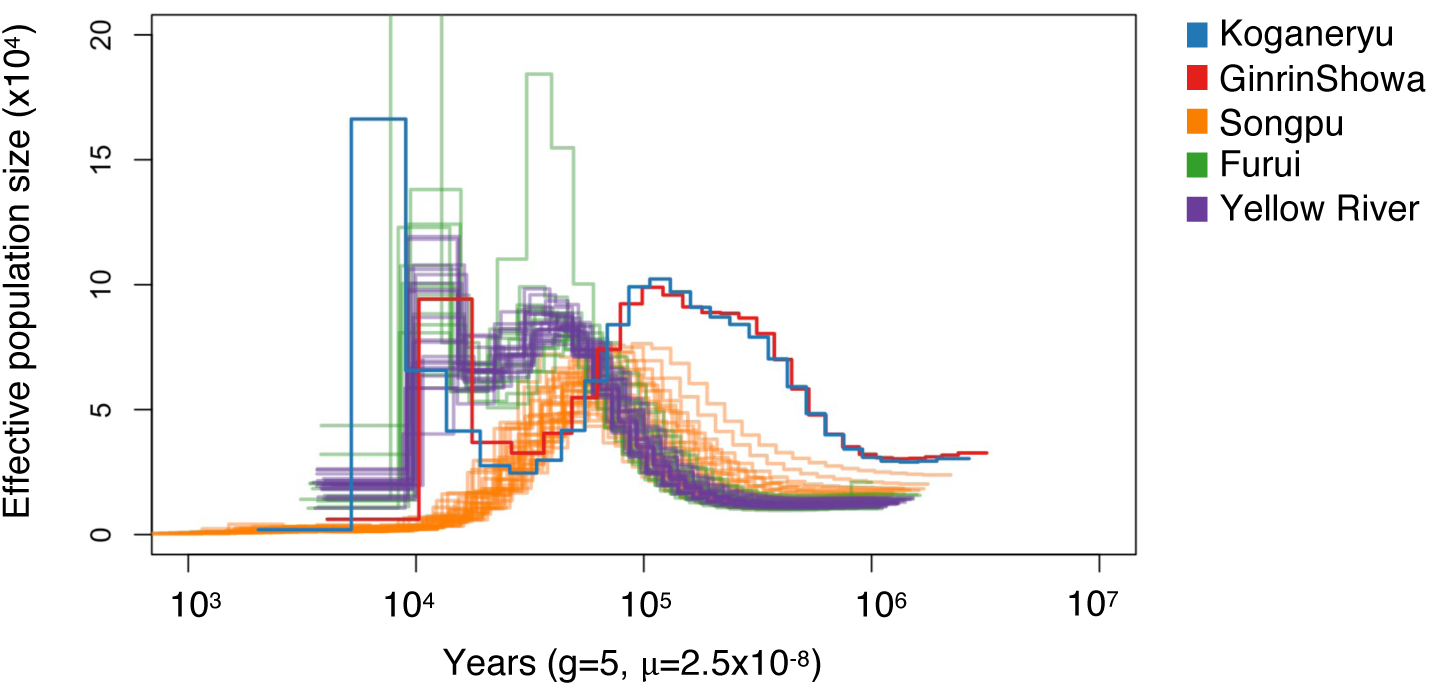
PSMC plot of Common carp strains and Nishikigoi strains. The fluctuation of effective population size over time among populations were plotted. X-axis and Y-axis indicate years and effective population size, respectively. Different colors indicate separate strains as shown in the legend.

**Supplementary Figure S3.**
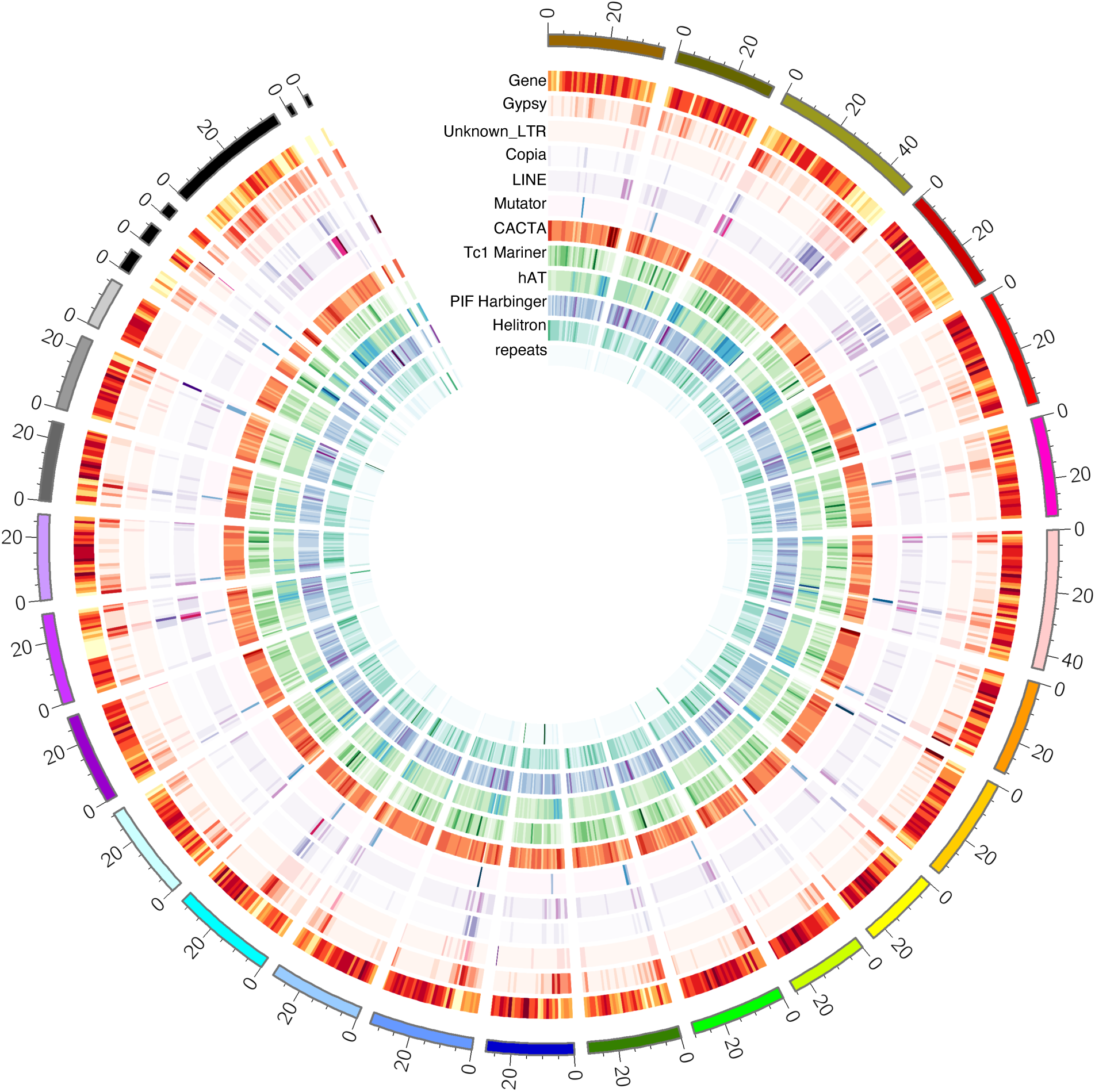
Distribution of TEs in the Nishikigoi genome. (A) Chromosomal distribution of genes, and transposons in B-genome of Nishikigoi genome. Heatmap indicates the density of the genes and the TE families in each 1Mb bin.

